# RNAi reveals a unique kinesin mediating chloroplast motility in the giant cytoplasm of *Bryopsis*, a coenocytic green alga

**DOI:** 10.1101/2025.05.19.654777

**Authors:** Harumi A. Ogawa, Kanta K. Ochiai, Maki Shirae-Kurabayashi, Moé Yamada, Gohta Goshima

## Abstract

RNA interference (RNAi) is a powerful tool for protein knockdown and is widely used in model animals and plants. Here, we applied this technique to *Bryopsis*, the green feather alga that develops a >10 cm coenocytic body in the wild and in laboratory culture. We mixed *in vitro*-transcribed double-stranded RNA (dsRNA) with extruded cytoplasm in the presence of polyethylene glycol or injected it directly into the cytoplasm, followed by thallus regeneration. After several days, we observed a reduction in the target gene transcript as well as expected phenotypes, indicating the effectiveness of RNAi. We prepared dsRNAs for the sole myosin and all 34 kinesin genes of the model *Bryopsis* strain, and performed RNAi and time-lapse microscopy to trace chloroplast movement. In addition to KCBP-type kinesins known to drive retrograde chloroplast transport in land plants, RNAi of a Bryopsidales-specific kinesin-14 (Kin14VIc) almost completely suppressed chloroplast motility. Cytoplasmic microtubules remained broadly aligned parallel to the main axis of the thallus following Kin14VIc RNAi. Purified Kin14VIc motor protein showed microtubule-gliding activity and, when artificially tetramerised, processive motility *in vitro* (∼250 nm/s), similar to plant KCBP. Thus, this study introduces a powerful gene loss-of-function tool in a coenocytic organism and identifies a uniquely evolved kinesin as a critical driver of chloroplast motility in the giant cytoplasm.

## Introduction

Marine siphonous (coenocytic) green algae present a fascinating model at the intersection of cell and developmental biology. These organisms consist of single, giant cells—extending over 30 cm in some species—that undergo complex morphogenesis and develop functionally distinct structures such as rhizoids, despite lacking cellular compartmentalisation (Graham *et al*. 2008). How macroscopic morphogenesis is achieved without cell division has long been a central question in algal biology (Dumais *et al*. 2000, Mine *et al*. 2008). However, the underlying mechanisms—particularly which genes are required for this phenomenon—remain poorly understood. This is largely due to the lack of model organisms with tools that allow large-scale gene screening for specific biological processes.

We have aimed to establish *Bryopsis*, a green feather alga, as an experimental model system for coenocytic cell biology. *Bryopsis* thalli grow over 10 cm tall and exhibit characteristic side branches and rhizoids (Mine, et al. 2008). However, nuclear division is not followed by septation, and there are no cell walls separating the many nuclei. These features are shared with other members of the order Bryopsidales, such as the edible *Codium* (green sea fingers) and *Caulerpa* (sea grapes). Uniquely, *Bryopsis* displays exceptional regenerative capacity: not only severed thalli, but also extruded cytoplasm, can rapidly reform membranes and regenerate into mature thalli (Kim *et al*. 2001, Tatewaki and Nagata 1970). This feature makes *Bryopsis* particularly amenable to laboratory culture; we have maintained both male and female strains in the lab for over four years. We recently assembled a high-quality genome for this species, providing a valuable resource for functional genomics (Ochiai *et al*. 2024).

In this study, we established two RNA interference (RNAi) protocols using synthesised double-stranded RNA (dsRNA), applied either to extruded cytoplasm or to severed thalli. Using this accessible and cost-effective system, we screened 35 cytoskeletal motor protein genes and identified a novel kinesin essential for chloroplast motility.

## Results and Discussion

### Development of an RNAi methodology based on extruded cytoplasm

We sought to develop a methodology to disrupt or silence genes of interest. We selected two nucleus-encoded genes, *ARC6* and *FtsZ*, which are required for chloroplast division in land plants (Pyke *et al*. 1994, Strepp *et al*. 1998). We expected that depletion of these gene products would result in the formation of giant chloroplasts in the *Bryopsis* cytoplasm, which would be readily detectable under a microscope.

After testing multiple approaches, we successfully developed an RNAi system in which dsRNA targets mRNA for degradation. We synthesised ∼1 kb dsRNAs targeting exon sequences of *ARC6* and the two *FtsZ* genes (*FtsZ1* and *FtsZ2*), and mixed them with extruded cytoplasm in the presence of polyethylene glycol (PEG) (Fig. 1A, B). After regeneration and thallus growth for seven or more days, enlarged chloroplasts were observed (Fig. 1C). The phenotype was frequent: 56% (n = 25) of thalli showed larger chloroplasts for *ARC6* RNAi, and 36% (n = 25) for *FtsZ1/2*. In contrast, control dsRNA with no homology to *Bryopsis* genes never produced abnormal chloroplasts (0%, n = 24). The RNAi effect lasted up to 14 days, after which normal-sized chloroplasts predominated, consistent with the transient nature of RNAi. Giant chloroplasts were also observed when alternative regions of the *FtsZ1/2* genes were targeted, ruling out off-target effects (Fig. 1B, C). Target gene knockdown was confirmed by quantitative RT-PCR (qRT-PCR) (Fig. 1D). These results show that RNAi-mediated knockdown works in *Bryopsis*, at least for the two chloroplast division genes, for a limited duration.

**Figure 1.**
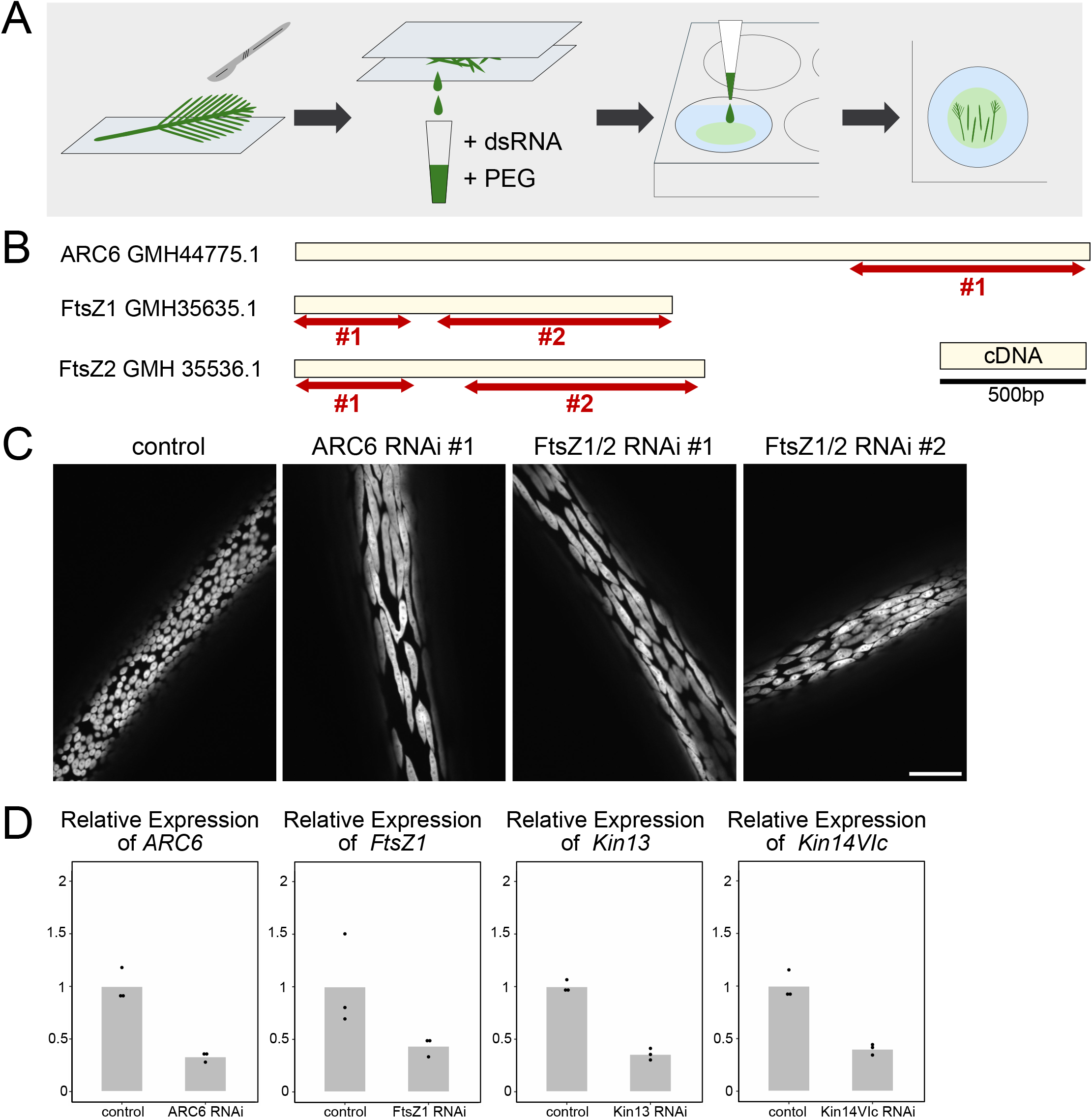
RNA interference (RNAi) using extruded cytoplasm in *Bryopsis*. (A) Procedure for PEG-based RNAi. The thallus was cut with a scalpel, sandwiched between two slide glasses, and crushed. The extruded cytoplasm was collected in a tube and mixed with the PEG/dsRNA solution. Protoplasts regenerate from the cytoplasmic solution in seawater. (B) RNAi constructs targeting the *ARC6* gene and two non-overlapping regions of *FtsZ1/2* genes. (C) Giant chloroplasts observed following *ARC6* or *FtsZ1/2* RNAi. Bar: 50 *δ*m. (D) qRT-PCR results showing knockdown of target genes seven days after dsRNA treatment. For *Kin13* and *Kin14VIc, ARC6* dsRNA was co-treated, and thalli with giant chloroplasts were selected. Experiments were performed in triplicate; bars represent mean relative expression levels.

### Development of an RNAi methodology based on cell perfusion

We developed an alternative method for RNA delivery. Tsubura (1990) reported in his PhD thesis that membrane closure following thallus severing is inhibited by EGTA. In his study, severed thalli were incubated in seawater containing EGTA and purified tubulin, and a portion of the cytoplasm was extracted using pipetting. Tubulin uptake into the cytoplasm was subsequently confirmed by electron microscopy (Tsubura 1990). We adapted this ‘cell perfusion’ method by adding *ARC6* dsRNA to the EGTA-containing seawater instead of tubulin (Fig. 2A). At day 4 post-treatment, giant chloroplasts were observed in 19 out of 20 perfused thalli, suggesting that RNAi is highly efficient with this method (Fig. 2B).

**Figure 2.**
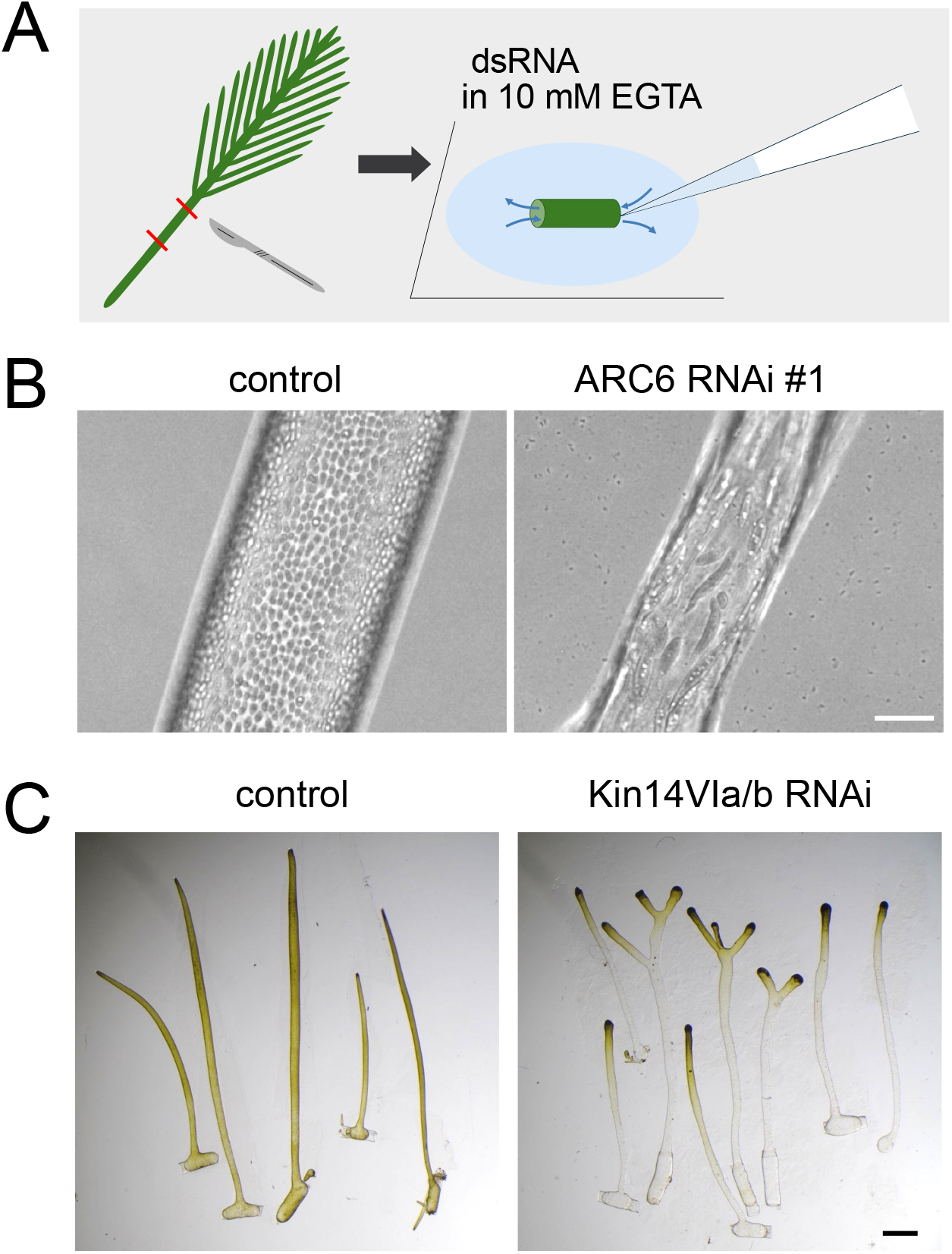
RNA interference (RNAi) using cell perfusion in *Bryopsis*. (A) Procedure for cell perfusion-based RNAi. (B) *ARC6* RNAi using the cell perfusion method. Bar: 50 *δ*m. (C) Apical accumulation of chloroplasts after double RNAi of *Kin14VIa* and *Kin14VIb*. Bar: 1 mm.

Because this method preserved the polarised structure of the thallus after RNAi, we next tested the effect of dsRNA targeting two KCBP kinesins (*Bryopsis* Kin14VIa and Kin14VIb). KCBP belongs to the kinesin-14VI subfamily and is known to mediate minus-end-directed transport of chloroplasts and other cargos, in a tug-of-war with the plus-end-directed motor KinARK in the moss *Physcomitrium patens*. In this species, knockout of all four KCBP paralogs results in chloroplast accumulation at the apex, where microtubule plus ends are enriched, due to the plus-end-directed motility of KinARK (Yamada *et al*. 2017, Yoshida *et al*. 2023, Yoshida *et al*. 2019). In *Bryopsis*, microtubules are similarly oriented with their plus ends towards the apex (Tsubura 1990), and required for bidirectional chloroplast motility (Ochiai, *et al*. 2024). We therefore hypothesised that KCBP knockdown in *Bryopsis* sp. KO-2023 would likewise lead to apical chloroplast accumulation. This was indeed observed: when dsRNAs targeting *Kin14VIa* and *Kin14VIb* were co-introduced using the cell perfusion method, chloroplasts accumulated at the apical tip of the thallus after six days (Fig. 2C).

Overall, the cell perfusion method induces phenotypes more rapidly and retains the polarised structure of the thallus. However, it is labour-intensive and challenging to scale up due to the precise pipetting steps required. In the present study, we employed both methods.

### RNAi screening identified a novel type of kinesin-14 as the motor protein required for chloroplast motility

To search for motor proteins responsible for chloroplast motility, we conducted an RNAi screen targeting all 34 kinesin genes in the *Bryopsis* genome. In addition, RNAi was performed on the sole myosin gene (Myosin XI), as a previous study suggested that actin might also play a role in chloroplast motility (Menzel and Schliwa 1986). We designed primer sets specific to each kinesin and myosin gene, amplified ∼800 bp cDNA fragments, and performed *in vitro* transcription to generate dsRNAs. Each dsRNA was mixed with extruded cytoplasm, which were allowed to regenerate into thalli. For two genes, mRNA reduction was validated by qRT-PCR, including one that showed no visible phenotype (*Kin13*), supporting the specificity and robustness of our RNAi system (Fig. 1D).

Between days 7 and 10 post-treatment, we performed time-lapse microscopy on 5– 12 regenerated thalli per treatment and tracked the movement of 20 chloroplasts per thallus (Fig. 3A). The screen did not identify Kin14VIa, Kin14VIb, or any putative plus-end-directed motors involved in chloroplast transport, possibly due to functional redundancy (i.e. phenotypes emerged only when dsRNAs targeting both Kin14VIa and Kin14VIb were introduced simultaneously). Strikingly, however, knockdown of the gene *GMH44105*.*1* caused a marked reduction in chloroplast motility in both directions, although chloroplasts remained uniformly distributed throughout the thallus (Fig. 3A, B). This phenotype was confirmed using two additional, non-overlapping dsRNA constructs (Fig. 3C, D). Importantly, microtubule organisation remained unaffected, indicating that the loss of motility was not a secondary effect of cytoskeletal disruption (Fig. 3E). The gene product was thus identified as a candidate motor for chloroplast motility in *Bryopsis*. Sequence analysis classified it as a member of the kinesin-14VI subfamily, which also includes KCBP, and we designated it Kin14VIc.

**Figure 3.**
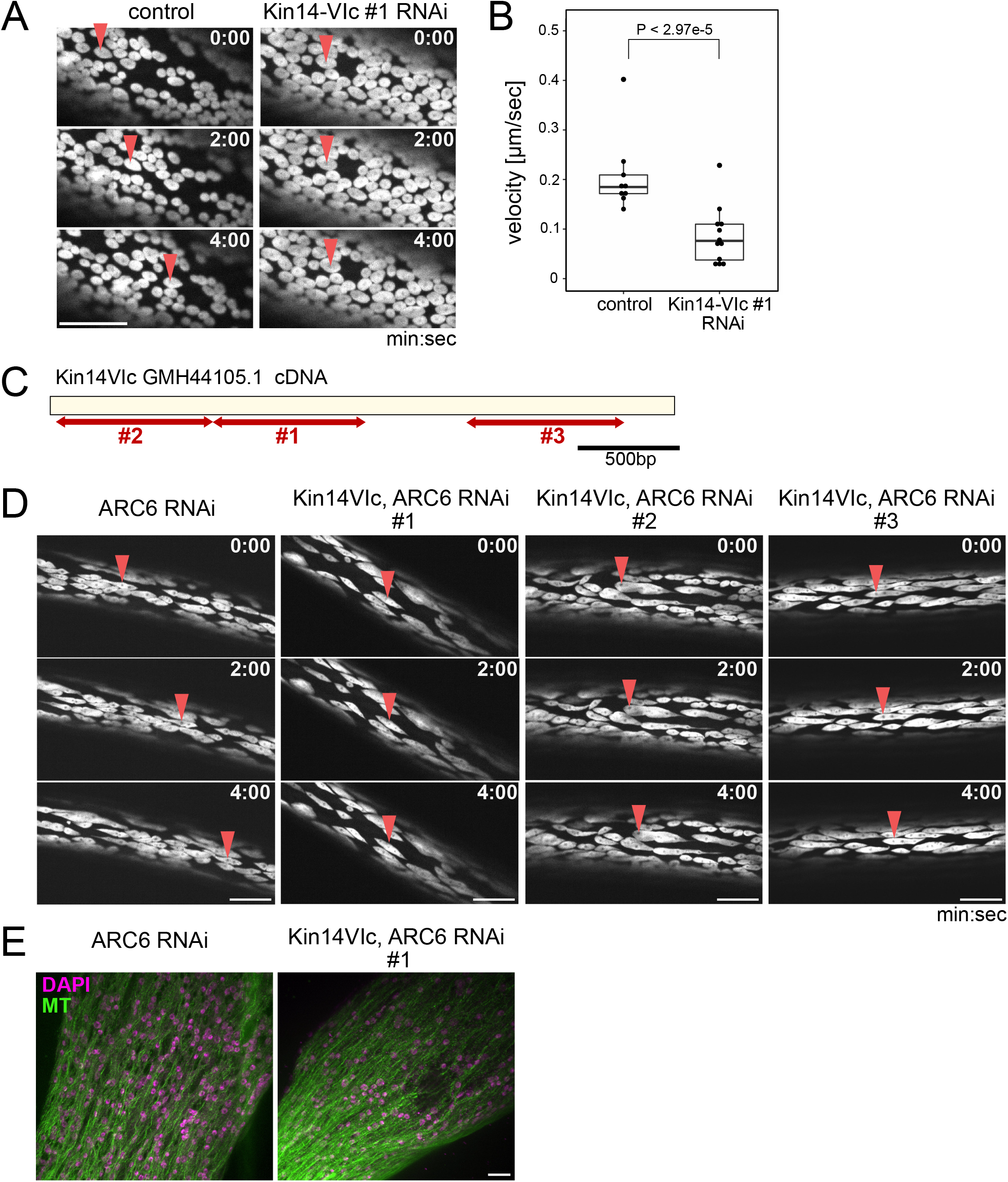
Kin14VIc is required for chloroplast motility. (A) Chloroplast motility. A motile (control) and a stationary (*Kin14VIc* RNAi-treated) chloroplast are indicated by red arrowheads. Bar: 50 *δ*m. (B) Chloroplast velocity. Nine (control) and twelve (*Kin14VIc* RNAi) thalli were analysed. Mean velocities: 208 ± 78 nm/s (control, ± SD, n = 180) and 86 ± 57 nm/s (*Kin14VIc* RNAi, ± SD, n = 240). Each dot indicates the mean velocity on the thallus from 9 or 12 samples, with the median and interquartile range indicated by black lines. P-value calculated using the Brunner–Munzel test. (C, D) Additional *Kin14VIc* dsRNA constructs also reduce chloroplast motility, indicated by red arrowheads. *ARC6* dsRNA was co-treated to ensure dsRNA incorporation in selected thalli. Bar: 50 *δ*m. (E) Immunostaining of microtubules in control and *Kin14VIc* RNAi-treated thalli. Images were processed using maximum z-projection of 1 *δ*m × 23 stacks (control) and 1 *δ*m × 28 stacks (*Kin14VIc* RNAi). Bar: 10 *δ*m.

Canonical KCBPs, including *Bryopsis* Kin14VIa and Kin14VIb, contain MYTH4 and FERM domains in the tail region, and the FERM domain is essential for chloroplast motility, likely serving as the cargo-binding domain (Shen *et al*. 2012, Yamada, et al. 2017, Yoshida, et al. 2019) (Fig. 4A). In contrast, Kin14VIc lacks MYTH4 and FERM domains and instead possesses a putative VHS (Vps27-Hrs-Stam) domain at its tail, a conserved protein-binding domain found across eukaryotes (Fig. 4A, B). VHS-containing Kin14VIc homologs are present in other Bryopsidales species (*Caulerpa lentillifera, Ostreobium quekettii*) but not in other green algae (Fig. 4C). However, BLAST searches using the VHS domain alone revealed highly similar sequences in other green algae such as *Chlamydomonas incerta* and *Coccomyxa* sp., although these are not part of kinesin proteins. These findings suggest that Kin14VIc likely emerged in the Bryopsidales lineage through the fusion of a KCBP-like motor domain with a VHS-containing, non-kinesin protein.

**Figure 4.**
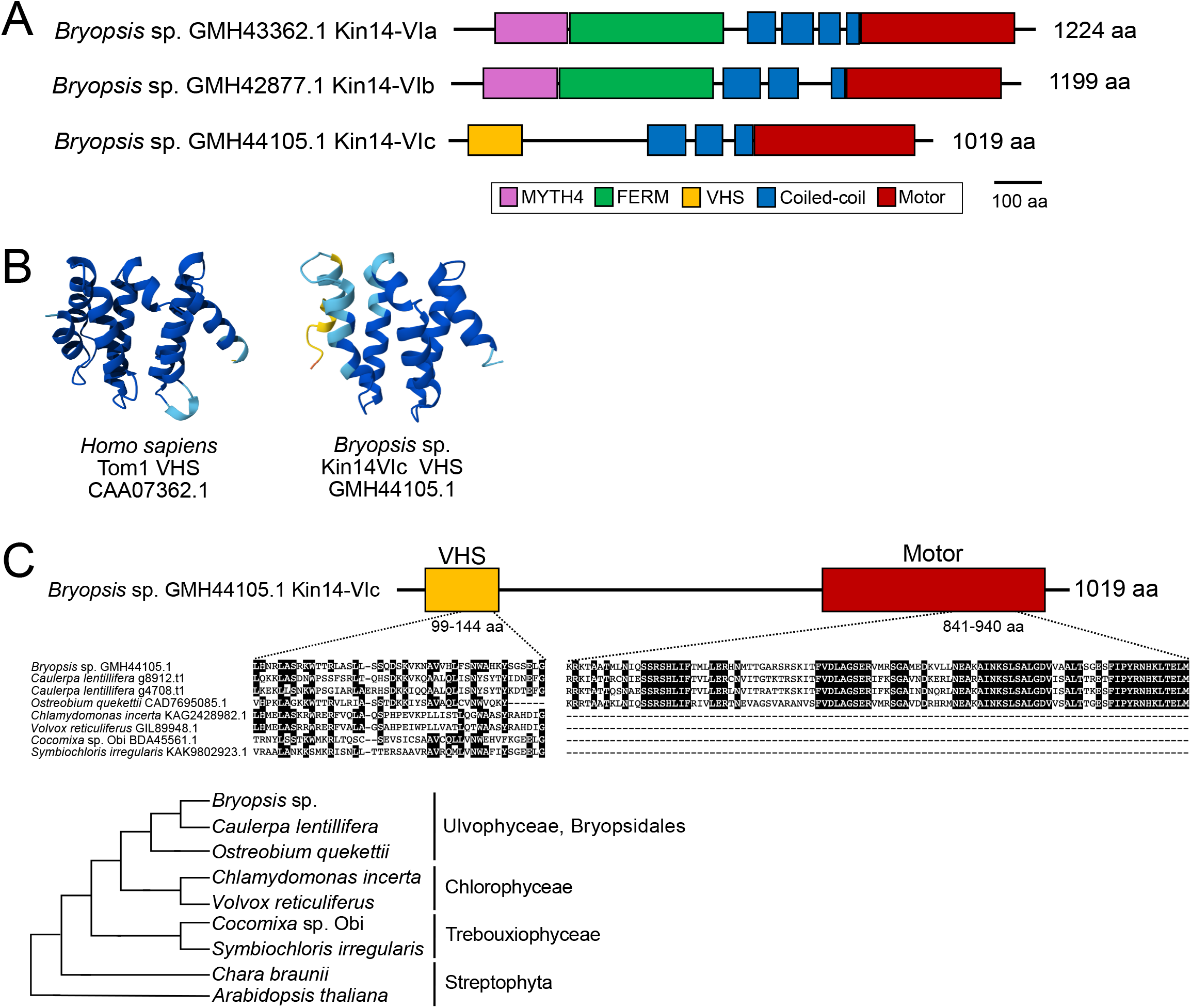
Kin14VIc evolved uniquely in Bryopsidales. (A) Schematic representation of the three *Kin14VI* subfamily proteins encoded by *Bryopsis*. (B) AlphaFold3-predicted structure of the *Bryopsis* VHS domain, compared with the known structure of the human Tom1 VHS domain. (C) Alignment of VHS domains and motor domains from green algal species. The phylogenetic relationship is illustrated at the bottom.

### Bryopsis Kin14VIc motor shows microtubule-based motility in vitro

We next investigated whether *Bryopsis* Kin14VIc exhibits microtubule-based motility, similar to KCBPs. We first purified a truncated version of the protein (474 amino acids), encompassing the motor domain and part of the coiled-coil region required for dimerisation (Fig. 5A, B). An EGFP tag was fused to the N-terminus for visualisation. In a conventional microtubule-gliding assay, Kin14VIc ΔN-motor immobilised on a glass surface translocated GMP-CPP-stabilised microtubules at a mean velocity of 43 ± 23 nm/s (SD, n = 205) (Fig. 5C, D). However, in a single-molecule motility assay, the dimeric motor bound to but rarely moved along surface-attached microtubules (Fig. 5E, left). This result is consistent with the known behaviour of most kinesin-14 family members, which are typically non-processive. Nevertheless, some kinesin-14 motors become processive when artificially oligomerised (Furuta *et al*. 2013, Jonsson *et al*. 2015). To test whether this also applies to Kin14VIc, we constructed a tetrameric version by fusing a GCN4 tetramerisation motif, following Jonsson et al. (2015). This motor construct (Kin14VIc GCN4^T^) retained microtubule-gliding activity (130 ± 35 nm/s, SD, n = 321) (Fig. 5C, D). In the single-molecule motility assay, the putative tetramer showed brighter fluorescence than the dimer, and displayed robust, unidirectional, processive motility (258 ± 72 nm/s, SD, n = 192; Fig. 5E, right; Fig. 5F). In some cases, the motor travelled distances longer than the length of the microtubule tracks themselves, indicating strong processivity. These findings suggest that multiple Kin14VIc motors can cooperate to transport cargo processively along microtubules.

**Figure 5.**
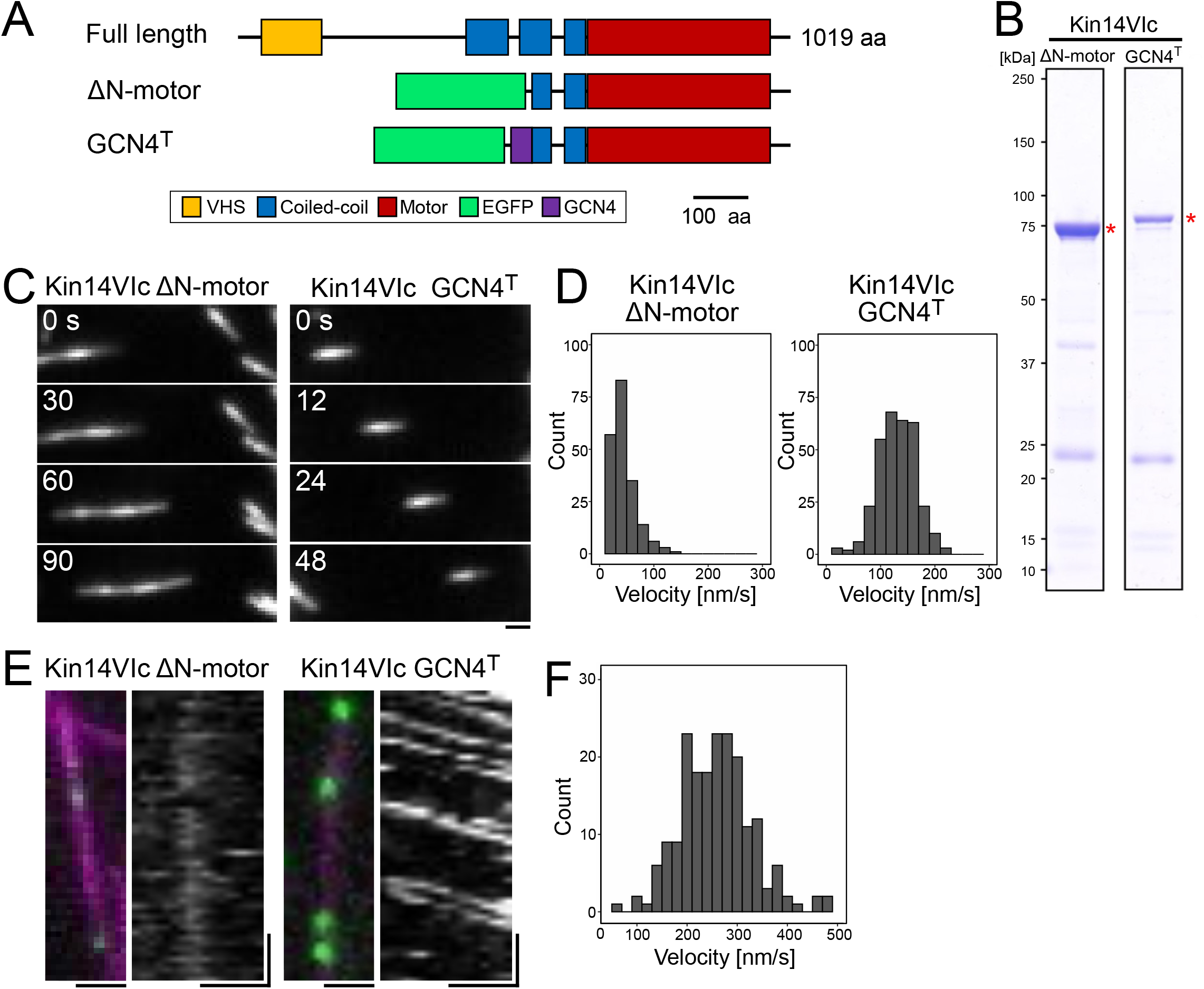
Kin14VIc exhibits microtubule-based motility *in vitro*. (A) Schematic of *Kin14VIc* constructs used in *in vitro* assays. The motor domain and part of the coiled-coil region (546–1019 a.a.) were used to construct the ΔN-motor and GCN4T. (B) SDS–PAGE gel images of purified proteins stained with Coomassie Brilliant Blue. The corresponding bands are indicated by asterisks. (C) Microtubule gliding activity of *Kin14VIc* proteins. Bar: 1 *δ*m. (D) Gliding velocity: *Kin14VIc* ΔN-motor, 43 ± 23 nm/s (mean ± SD, n = 205); *Kin14VIc* GCN4T, 130 ± 35 nm/s (n = 321). Results pooled from three independent experiments. (E) Motility of *Kin14VIc* along microtubules (magenta: microtubules; green: *Kin14VIc*). Corresponding kymographs are also shown. Bars: (left) 1 *δ*m; (right) horizontal 1 *δ*m, vertical 10 s. (F) Velocity of processive movement of *Kin14VIc* GCN4T: 258 ± 70 nm/s (mean ± SD, n = 192).

Altogether, our results suggest that *Bryopsis* Kin14VIc is a microtubule-based motor critical to chloroplast motility, potentially operating via a mechanism distinct from canonical KCBPs. However, the precise manner in which Kin14VIc contributes to chloroplast motility remains unclear.

### RNAi as a powerful tool to elucidate gene function in *Bryopsis*

To our knowledge, this is the first method enabling specific gene perturbation in Bryopsidales. Our approach, using 500–1000 bp double-stranded RNA (dsRNA), builds on established protocols from model animal systems such as *C. elegans* and *Drosophila* S2 cells (Clemens *et al*. 2000, Fire *et al*. 1998), while PEG is commonly used for transformation of walled organisms (Moreno *et al*. 1991, Oertel *et al*. 2015, Yamada *et al*. 2016). The cell perfusion method was a modification of the protein injection technique described by Tsubura (1990). These methods are simple, scalable, and cost-effective, requiring neither siRNA synthesis, specialised transfection reagents, nor plasmid construction for hairpin RNA expression.

As a proof of principle, we applied the RNAi system to 35 motor genes and identified an unexpected kinesin required for chloroplast motility. We also demonstrated that simultaneous knockdown of two genes—such as *ARC6* and *Kin14VIc*, or *Kin14VIa* and *Kin14VIb*—is feasible simply by mixing the respective dsRNAs. While the PEG-based approach may be limited to *Bryopsis*, which uniquely regenerates from extruded cytoplasm, the alternative cell perfusion method may be more broadly applicable across Bryopsidales, as other species also exhibit high regenerative capacity following severing (Shirae-Kurabayashi *et al*. 2022).

However, RNAi inherently produces incomplete depletion of target proteins, and our system is no exception. Thus, negative results from RNAi cannot be considered definitive. In this study, we cannot exclude the possibility that other motor proteins contribute to chloroplast motility but remained undetected due to residual protein levels after knockdown. To achieve complete loss of function, the development of a gene knockout technique is essential. A promising strategy would be to implement CRISPR/Cas9-based genome editing, potentially in combination with either the cytoplasm/PEG or cell perfusion delivery methods.

## Materials and methods

### *Bryopsis* culture

We followed the protocol described in Ochiai (2024). Briefly, haploid *Bryopsis* thalli were cultured daily at 15 °C under a 16 h light / 8 h dark cycle (90 μmol m^−2^ s^−1^) in filtered ocean surface water (salinity 2.8–3.4%). The seawater was filtered using a 0.22-μm Millipore Stericup, autoclaved, and supplemented with Daigo’s IMK medium (252 mg/L, Shiotani M.S.). Cultures were maintained by severing the thalli and transferring them to fresh medium.

### RNA interference

[PEG method] Double-stranded RNA (dsRNA) was synthesised following the method used for RNAi in the *Drosophila* S2 cell line (Bettencourt-Dias and Goshima 2009). Briefly, templates for *in vitro* transcription were generated by PCR using the primers listed in Table S1, with T7 promoter sequences added to the 5′ end of each primer. The synthesised dsRNA (10 μg) was mixed with 40 μL of PEG 1540 (1 g dissolved in 1 mL distilled water; KANTO CHEMICAL CO., INC., Tokyo, Japan). The *Bryopsis* thallus was cut with a scalpel and crushed between two glass slides. The extruded cytoplasm was collected into a tube, and 40 μL of it was added to the PEG/RNA mixture. The solution was mixed gently by tapping for 5 min, then incubated at room temperature for another 5 min. Autoclaved seawater (1 mL) was added, followed by a 10 min incubation. The upper, transparent portion of the seawater (not enriched in cytoplasm) was discarded. The remaining cytoplasm-containing solution was mixed by pipetting and transferred into 8 mL of autoclaved seawater in a 6-well plate.

[Cell perfusion method] We modified the protocol described in Tsubura (1990). A 100 μL droplet of Daigo’s artificial seawater (36 g/L; Shiotani M.S.) containing 10 mM EGTA was prepared, and a 1-cm fragment from the main axis of *Bryopsis* was placed in the droplet. This fragment was then cut into smaller pieces (0.2–0.5 cm) using a razor blade, and a pair of forceps was used to form tubular fragments with both ends open. A Pasteur pipette, heat-drawn using an alcohol lamp, was inserted into the hollow fragment, and the solution was perfused by alternately aspirating and injecting it 2–3 times. For RNAi, 10 μg of dsRNA was added to the 100 μL droplet before perfusion.

### Quantitative RT-PCR

At days 5, 7, and 9 after RNAi treatment, total RNA was extracted from ∼80 thalli using the RNeasy Plant Mini Kit (Qiagen), with DNase treatment included. For kinesin RNAi samples collected on days 7 and 9, dsRNAs against both kinesin and *ARC6* were co-introduced, and thalli with visibly enlarged chloroplasts were selectively harvested for RNA extraction—thus enriching for successfully treated samples. This selection could not be applied to day-5 samples, as the chloroplast phenotype was not yet penetrant at that early stage. Extracted RNAs were reverse transcribed into cDNA using Takara PrimeScript II. Quantitative RT-PCR (qRT-PCR) was conducted in triplicate using PowerUp™ SYBR™ Green Master Mix and the 7500 Real-Time PCR System (Model 7500; Applied Biosystems). Expression levels were normalised to the *EF1α* gene. Primers used for qRT-PCR are listed in Table S2. While results from day 7 are shown as representative, in some cases, RNA reduction was more pronounced at day 5.

### Immunostaining

Immunostaining of microtubules was performed following the protocol described in Ochiai et al. (2024). Briefly, thalli 4 days after dsRNA treatment via the cell perfusion method were fixed in 4% paraformaldehyde in modified PHEM buffer (60 mM PIPES, 25 mM HEPES, 0.5 M NaCl, 10 mM EGTA, 2 mM MgCl_2_; pH 6.9) for 1 h at 25 °C. Samples were then permeabilised with 1% Triton X-100 in PBS for 1 h at 25 °C. After two washes with PBST (0.1% Triton X-100 in PBS), specimens were incubated in blocking solution (1% BSA in PBST) for 1 h at 25 °C, followed by overnight incubation at 4 °C with rotation in primary antibody (rat anti-α-tubulin [YOL1/34, MCA78G, Bio-Rad], 1:1000). Samples were washed three times with PBST and incubated overnight at 4 °C with rotation in secondary antibody (anti-rat, Jackson ImmunoResearch, 712-165-153, 1:1000) and DAPI (final concentration 1 μg/mL). After two final PBST washes, specimens were mounted on glass slides using Fluoromount™ (Diagnostic BioSystems).

### Microscopy

Microscopy was conducted basically as described in Ochiai et al. (2024). The entire thallus was imaged with Nikon SMZ800N stereo microscope with Plan Apo 19/WF lens and NY1S-EA camera (SONY). Transmission light images of chloroplasts were acquired using a BZ-X710 fluorescence microscope (Keyence) equipped with an S Plan Fluor 20× 0.45 NA lens (Nikon). Fluorescence images of DNA (DAPI), chloroplasts, and microtubules were captured using a Nikon Ti2 inverted microscope fitted with a CSU-10 or CSU-W1 spinning-disc confocal scanner unit (Yokogawa), a Zyla CMOS camera (Andor), and laser lines at 637, 561, 488, and 405 nm. A 40× 0.95 NA or 100× 1.45 NA lens was used for imaging live or fixed cells, respectively. For quantifying chloroplast motility, a 35-mm glass-bottom dish was prepared with a piece of kitchen garbage net (∼10 × 20 mm) affixed using double-sided tape. Following cytoplasmic extrusion, a 3-week-old thallus and a coverslip were placed on the net, and 1 mL of autoclaved seawater was added. The net served to immobilise the thallus during imaging. Autofluorescent chloroplasts were imaged every 10 s using the spinning-disc confocal microscope and a 40× 0.95 NA lens. The unidirectional motility rates of randomly selected chloroplasts were measured manually from kymographs generated in Fiji.

### Protein purification

Sequences of the Kin14VIc dimer and tetramer constructs are provided in the Supplementary Files (SnapGene format). For the tetramer, a GCN4 tetramerisation motif was added following a previous study (Jonsson, et al. 2015). Proteins were expressed in *E. coli* SolBL21 cells using 0.2 mM IPTG at 15 °C for 20 h. Harvested cells were washed with PBS, and the bacterial pellet was lysed using an Advanced Digital Sonifier D450 (Branson) in lysis buffer (25 mM MOPS [pH 7.0], 250 mM KCl, 2 mM MgCl_2_, 1 mM EGTA, 20 mM imidazole, 0.1 mM ATP), supplemented with 5 mM β-mercaptoethanol, 500 U benzonase, and protease inhibitors (1 mM PMSF and a 5000× peptide inhibitor cocktail: 5 mg/mL each of aprotinin, chymostatin, leupeptin, and pepstatin A). After centrifugation, the clarified lysate was incubated with nickel-NTA agarose beads for 1.5 h at 4 °C. The beads were washed five times with the lysis buffer, and proteins were eluted with 1000 μL of elution buffer (25 mM MOPS [pH 7.0], 75 mM KCl, 2 mM MgCl_2_, 1 mM EGTA, 200 mM imidazole, 0.2 mM ATP) supplemented with 5 mM β-mercaptoethanol and 20% (w/v) sucrose. Freshly purified proteins were used immediately in assays. Protein concentration was determined by Coomassie staining of SDS–PAGE gels alongside a BSA standard.

### *In vitro* assay

Microtubule polymerisation was performed using a biotinylated tubulin mixture composed of 10% biotinylated pig tubulin and 30% Alexa Fluor 568-labelled pig tubulin, at a total concentration of 100 μM in the presence of 0.5 mM GMPCPP, prepared in MRB80 buffer (80 mM PIPES-KOH, pH 6.8, 1 mM EGTA, 4 mM MgCl_2_). The mixture was incubated at 37 °C for 40 min to induce polymerisation. For the gliding assay using purified EGFP-Kin14VIc, 1× Standard Assay Buffer (SAB: 25 mM MOPS [pH 75 mM KCl, 2 mM MgCl_2_, 1 mM EGTA) was used. Briefly, 7 μL of purified protein was introduced into a flow chamber made with a silanised coverslip and incubated at room temperature for 5 min in the dark. A 10 μL reaction mixture (1× SAB, 0.1% methylcellulose, 50 mM glucose, 0.5 μg/μL κ-casein, GMPCPP-stabilised microtubule seeds, oxygen scavenger system, 1 mM ATP) was then introduced into the chamber, which was sealed with candle wax. Alexa Fluor 568-labelled GMPCPP-stabilised microtubules were imaged by TIRF microscopy every 3 s for 10 min at 23–25 °C. For the single-molecule motility assay with purified EGFP-Kin14VIc, the same SAB buffer was used. A silanised coverslip was coated with an anti-biotin antibody, followed by introduction of SAB containing 1% pluronic acid. After one wash with SAB, GMPCPP-stabilised microtubules were introduced in the presence of 20 μM Taxol and incubated for 5 min. The chamber was washed once with SAB containing 20 μM Taxol, followed by introduction of 10 μL reaction mixture (1× SAB, 0.1% methylcellulose, 50 mM glucose, 0.5 μg/μL κ-casein, 20–200 pM Kin14VIc, 75 mM KCl, oxygen scavenger system, 1 mM ATP, 20 μM taxol). The chamber was then sealed with candle wax. TIRF imaging was performed every 1 s for 2 min at 23–25 °C.

## Supporting information

Plasmid seq

## Acknowledgements

This work was supported by the Japan Society for the Promotion of Science KAKENHI grants to G.G. (22K19308 and 23K23907), the Shiodamari Foundation to K.K.O., and the Japan Science and Technology Agency ACT-X grant to M.Y. (JPMJAX242J). The authors declare no conflict of interest.

## Author contributions

H.A.O. and G.G. designed research; H.A.O., K.K.O. and M.S-K. developed methodology; H.A.O. and M.Y. performed experiments and analysed the data; G.G. wrote the paper.

## Movie legends

**Movie 1. Kin14VIc-dependent chloroplast motility**

Time-lapse movie of chloroplasts using spinning-disc confocal microscopy. Bar: 20 μm.

**Movie 2. Microtubule-gliding activity of Kin14VIc motors *in vitro***

Microtubule gliding by EGFP-Kin14VIc ΔN-motor (dimeric) and EGFP-Kin14VIc GCN4^T^ (tetrameric) constructs. Images were acquired with total internal reflection fluorescence (TIRF) microscopy. Bar: 1 μm.

**Movie 3. Artificially tetramerised Kin14VIc motor shows processive motility *in vitro*** Tetrameric EGFP-Kin14VIc GCN4^T^ (green) and microtubules (magenta) were imaged with total internal reflection fluorescence (TIRF) microscopy. Bar: 1 μm.

## Supplementary tables

**Table S1:**
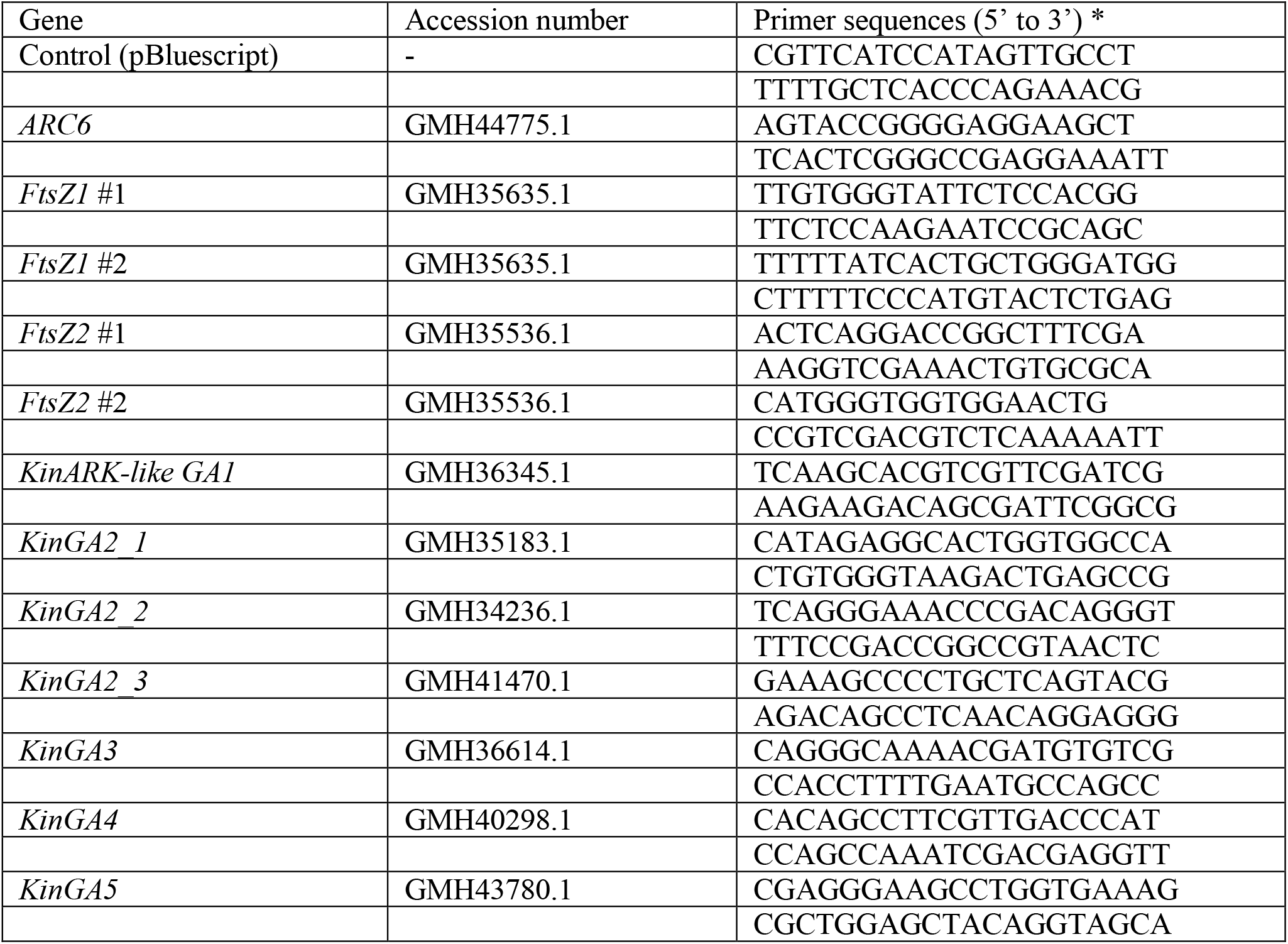
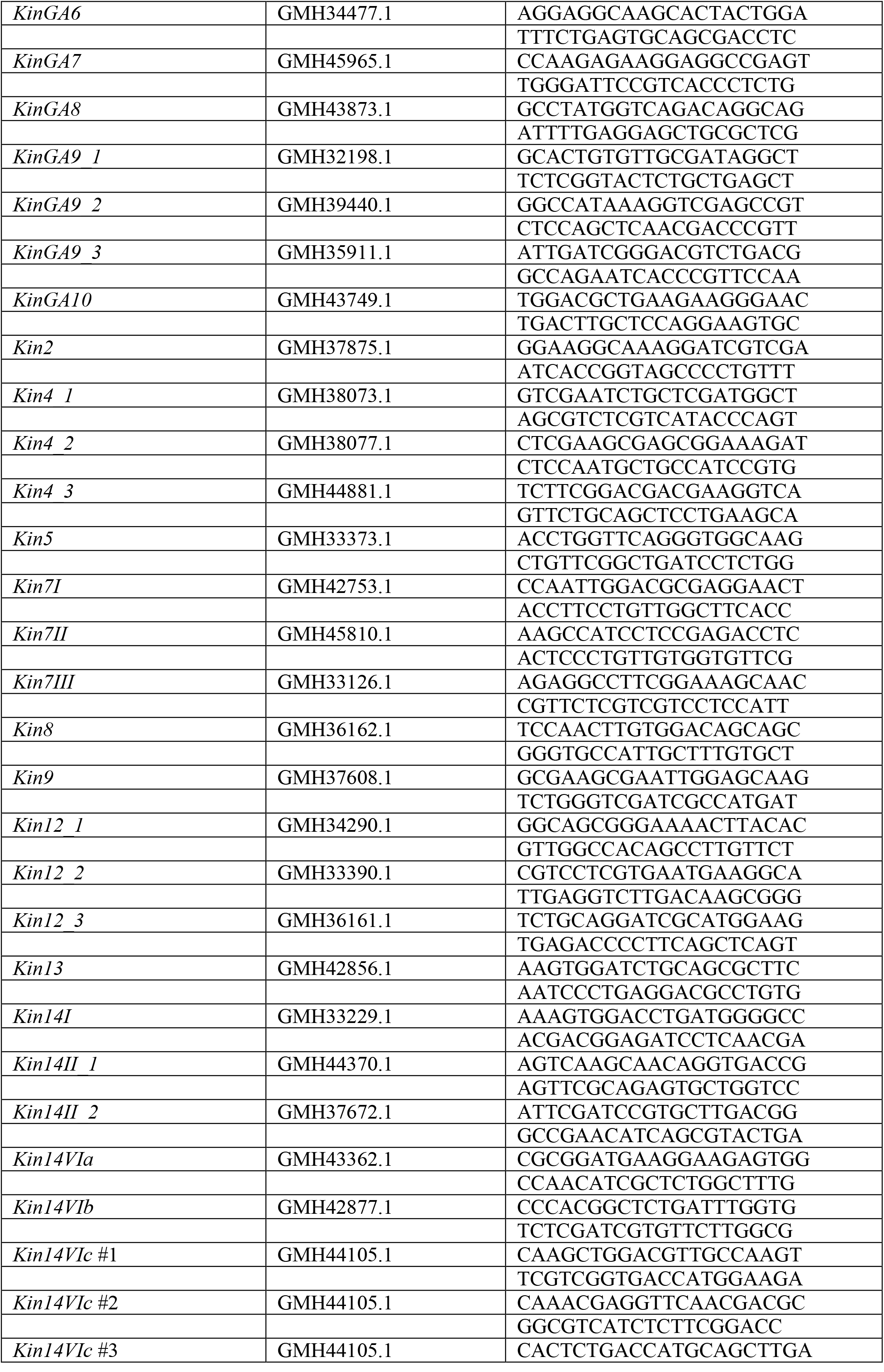
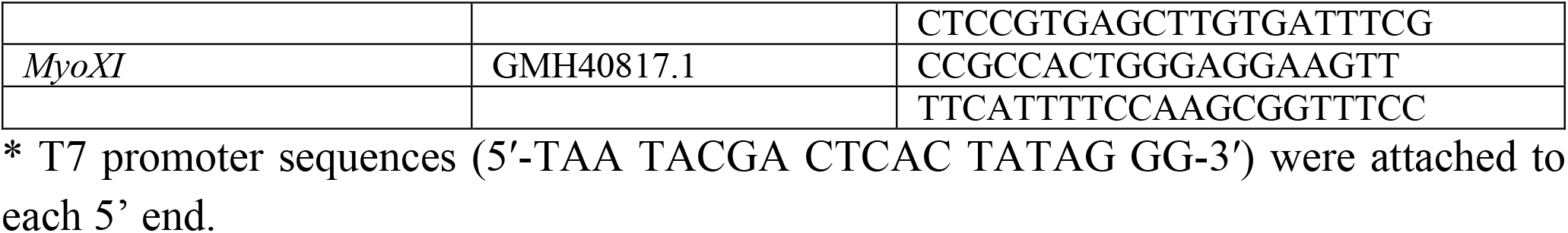
Primers for RNAi.

**Table S2.**
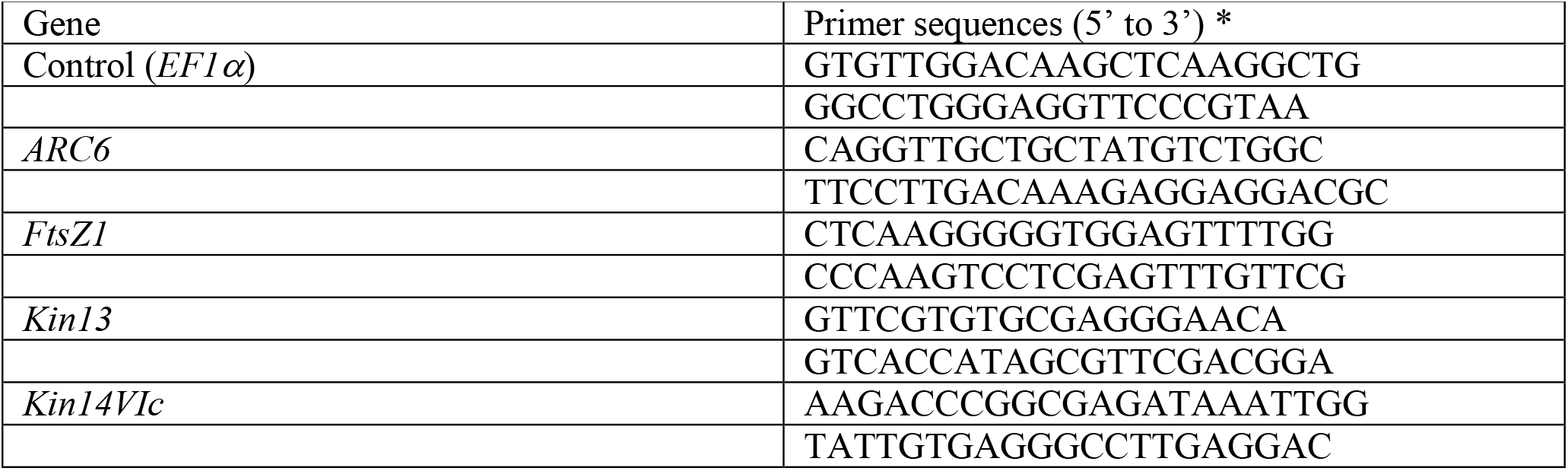
Primers for qRT-PCR.

